# Identification and characterization of phlorizin transporter from *Arabidopsis thaliana* and its application for phlorizin production in *Saccharomyces cerevisiae*

**DOI:** 10.1101/2020.08.14.248047

**Authors:** Zeinu Mussa Belew, Christoph Crocoll, Iben Møller-Hansen, Michael Naesby, Irina Borodina, Hussam Hassan Nour-Eldin

## Abstract

Bioengineering aimed at producing complex and valuable plant specialized metabolites in microbial hosts requires efficient uptake of precursor molecules and export of final products to alleviate toxicity and feedback inhibition. Plant genomes encode a vast repository of transporters of specialized metabolites that— due to lack of molecular knowledge—remains largely unexplored in bioengineering. Using phlorizin as a case study—an anti-diabetic and anti-cancerous flavonoid from apple—we demonstrate that brute-force functional screening of plant transporter libraries in *Xenopus* oocytes is a viable approach to identify transporters for bioengineering. By screening 600 *Arabidopsis* transporters, we identified and characterized <u>pu</u>rine <u>p</u>ermease 8 (AtPUP8) as a bidirectional phlorizin transporter. Functional expression in the plasma membrane of a phlorizin-producing yeast strain increased phlorizin titer by more than 80 %. This study provides a generic approach for identifying plant exporters of specialized metabolites and demonstrates the potential of transport-engineering for improving yield in bioengineering approaches.

## 1. Introduction

Plants synthesize a vast number of specialized small molecules that have pharmaceutical and nutraceutical values. These molecules are usually produced in minute amounts in the natural plant host, wherefrom extraction may be costly, insufficient and time-consuming (Liu et al., 2017; Lv et al., 2016). As a promising alternative, bioengineering in microorganisms such as *Saccharomyces cerevisiae* and *Escherichia coli* has emerged as a sustainable and cost-effective means for large-scale production of high-value plant natural products (Keasling, 2012; Krivoruchko and Nielsen, 2015; Suzuki et al., 2014).

For any bioengineering approach, transport proteins play critical roles in uptake of nutrients, cofactors and precursors and also for secretion of final products to alleviate toxicity and feedback inhibition that may hamper yield (Jones et al., 2015; Lee et al., 2012; Liu et al., 2017; Shu and Liao, 2002).

Transporters native to the producing microbial cell factory (endogenous transporters) or heterologous transporters obtained from other organisms can be utilized in transport-engineering approaches aimed at improving these transport processes. Most studies implementing transport-engineering have so far targeted microbial transporters. For example, tolerance of *E. coli* was improved by applying directed evolution to enhance the export efficiency of AcrB towards toxic biofuels and plastic precursors (Fisher et al., 2014; Foo and Leong, 2013). In other examples, overexpression of YddG, MsbA and YadH improved secretion and thereby production of various amino acids, biofuels and plant natural products such as anthocyanins in *E. coli* (Doroshenko et al., 2007; Doshi et al., 2013; Lim et al., 2015).

Given the enormous diversity of chemical structures in plant specialized metabolism, the field of transport-engineering will require access to a large repository of transporters. The general participation of separate organs and/or cells in the synthesis and accumulation of plant specialized metabolites indicates that plant genomes encode a plethora of specialized metabolite transporters (Shitan, 2016; Yazaki, 2005). However, in the context of bioengineering, plant transporters remain largely unexplored.

In this study, we seek to address two challenges; i) how to identify plant transporters relevant for a given bioengineering approach and ii) demonstrating the potential of using plant transporters to improve production in a given bioengineering approach.

As a case study, we focus on phlorizin, a dihydrochalcone *O*-glucoside predominantly found in apple (*Malus sp*.), which is comprised of a glucose moiety and two aromatic rings joined by an alkyl spacer. Phlorizin is medicinally important as it decreases blood glucose levels, has been used as both a remedy and a tool to study renal physiology for nearly two centuries and has been used as a blueprint to develop several commercialized anti-diabetic drugs (Ehrenkranz et al., 2005; Kramer and Zinman, 2019; Meng et al., 2008; Nomura et al., 2010; Scheen, 2015).

Recently, a pathway consisting of seven enzymes was engineered into *S. cerevisiae* for *de novo* production of phlorizin (Eichenberger et al., 2017). The titer is low for industrial production indicating that optimization of the biosynthesis pathway is possible, for example by increasing cytoplasmic malonyl-CoA, altering gene copy number, enzyme origins, and culture conditions (Galanie and Smolke, 2015; Li et al., 2015; Rodriguez et al., 2015). Although precursor supply (such as malonyl-CoA, a) is a major bottleneck for microbial production of flavonoids in general and phlorizin in particular (Delmulle et al., 2018; Eichenberger et al., 2017), product toxicity, negative feedback inhibition by intermediates (such as cinnamic acid and p-coumaric acid) on early biosynthetic enzymes (Blount et al., 2000; Lam et al., 2008; Sarma and Sharma, 1999) and by-product UDP inhibition on the last biosynthetic enzyme UGT (Zhang et al., 2016) could hamper overall flux into the phlorizin pathway.

Phlorizin is a polar compound that requires a membrane-bound transporter protein to traverse membranes. In this study, we explore whether titer can be increased by expressing an efficient phlorizin exporter to improve product secretion from the phlorizin producing yeast strain. To test this hypothesis, we faced a typical knowledge gap in plant specialized metabolism, which is that as of date, no phlorizin transporters have been identified.

We have developed a functional genomics approach wherein we build and functionally screen sequence-indexed full-length transporter cDNA libraries in *Xenopus* oocytes for activity towards target compounds (Nour-Eldin et al., 2006). Here, we screen a library consisting of 600 transporters from *Arabidopsis* and identify the first reported phlorizin transporter. Extensive biophysical characterization in *Xenopus* oocytes reveals a passive, medium-affinity, proton gradient-independent, bidirectional transport activity, which when introduced into a phlorizin producing yeast strain increased titer significantly. The transporter belongs to the purine permease family (PUP) present in all plants, which here has its substrate spectrum expanded to include a novel class of specialized metabolites. This study provides a generic approach for identifying plant exporters of specialized metabolites and demonstrates the potential of transport-engineering for improving yield in bioengineering of plant specialized metabolites.

## 2. Materials and Methods

### 2.1 Transporter cDNA library and in vitro transcription

The transporter library screened in this study contains 600 full-length cDNAs encoding *Arabidopsis* membrane proteins (unpublished). The CDSs are in *Xenopus* expression vectors, either in pNB1u or in pOO2-GW vector.

Linear DNA templates for *in vitro* transcription were generated from pNB1u or pOO2-GW plasmid by PCR using Phusion High-Fidelity DNA Polymerase (NEB), according to the manufacturer’s instructions. Primer pairs of HHN49 and HHN50 were used to generate templates from pNB1u plasmid (Wulff et al., 2019), whereas Bolar051 FW1 and Bolar051 RV2 pairs were used to PCR amplify from pOO2-GW plasmid (Larsen et al., 2017a) (Supplementary Table S1). PCR products were purified using the QIAquick PCR Purification Kits (Qiagen), according to the manufacturer’s instructions. Capped cRNA was *in vitro* synthesized using the mMessage mMachine T7 Kit (Ambion) following the manufacturer’s instructions. Concentration of the synthesized cRNA of each transporter gene was normalized to 800 ng/μl.

### 2.2 Expression in Xenopus oocytes

Defolliculated *X. laevis* oocytes, stage V-VI, were purchased from Ecocyte Bioscience (Germany). Oocytes were injected with 50.6 nl cRNA using a Drummond NANOJECT II (Drummond scientific company, Broomall Pennsylvania). For the transporter library screening, cRNA of 8 genes were pooled (final concentration of 100 ng/μl of each cRNA) and injected into oocytes. For single gene expression, ~250 ng/μl of cRNA was used for injection. Injected oocytes were incubated for 3 days at 16 °C in HEPES-based kulori buffer (90 mM NaCl, 1 mM KCl, 1 mM MgCl_2_, 1 mM CaCl_2_, 5 mM HEPES pH 7.4) supplemented with gentamycin (100 μg/mL). For non-expressing negative control oocytes (mock-injected control), 50.6 nl nuclease-free water (Ambion) was injected instead of cRNA.

### 2.3 Transport assays in Xenopus oocytes

Uptake assay in *X. laevis* oocytes was performed essentially as described previously (Jørgensen et al., 2017a), with some modifications. Three days after cRNA injection, oocytes were pre-incubated for 5 min in 5 ml MES-based kulori buffer (90 mM NaCl, 1 mM KCl, 1 mM MgCl_2_, 1 mM CaCl_2_, 5 mM MES pH 5.0), then incubated in 0.5 ml MES-based kulori buffer (pH 5.0) containing a mixture of phlorizin (0.5 mM) and 4MTB (0.1 mM) for 1 h. Oocytes were washed 3 times in kulori buffer (>20 ml) and homogenized with 50 % methanol (containing internal standard) and stored at −20 °C overnight. Subsequently, oocyte extracts were spun down at 10000 x *g* for 10 min at 4 °C, supernatants were transferred to new Eppendorf tubes and spun down at 10000 x *g* for 10 min at 4 °C. The supernatant was diluted with water and filtered through a 0.22 μm filter plate (MSGVN2250, Merck Millipore) and analyzed by LC-MS/MS as described below.

For export assay: 3 days after cRNA injection, oocytes were injected with 23 nl of 21.7 mM phlorizin or 4MTB (to obtain an initial internal concentration of ~0.5 mM) before a 5 min wait to reseal the hole (caused by the injection) before washing 3 times in kulori buffer (pH 7.4). Subsequently, some oocytes were harvested for T = 0 samples and the remaining oocytes were incubated in kulori buffer (pH 7.4) in a 96-well U-bottom microtiter plate (Greiner Bio-One) (3 oocytes in a well containing 200 μl buffer). Oocytes and the corresponding external medium were harvested at different time points over 22 h. The harvested oocytes were washed 3 times in kulori buffer (pH 7.4) and homogenized with 50 % methanol. 10 μl of the external medium was sampled, homogenized with 50 % methanol, and then treated like the oocyte samples. The extraction and filtration through a 0.22 μm filter plate were done as mentioned above. Finally, samples were analyzed by LC-MS/MS as described below. For both import and export assays, oocyte concentration of phlorizin and 4MTB was calculated based on an estimated cytosolic oocyte volume of 1 μl (Jørgensen et al., 2015).

### 2.4 Yeast strains, plasmids, and cloning

*S. cerevisiae* strain BG (*MATα hoΔ0 his3Δ0 leu2Δ0 ura3Δ0 cat5Δ0::CAT5(I91M) mip1Δ0::MIP1(A661T) gal2Δ0::GAL2 sal1Δ0::SAL1*) obtained from a previous study (Eichenberger et al., 2017) was used for all yeast experiments in this study. Precursor phloretin-producing platform strain DBR2 (in the BG background strain) was obtained from a previous study (Eichenberger et al., 2017). Strain DBR2 is based on plasmid (URA3 as a selection marker) expressing six genes (*AtPAL2, AmC4H, ScCPR1, At4CL2, HaCHS* and *ScTSC13*). Other yeast strains developed using BG as background strain are listed in Supplementary Table S2. Expression cassette plasmid backbone pEVE2176 carrying the *Pyrus communis* UDP-dependent-glycosyltransferase (UGT) gene *PcUGT88F2* was kindly provided by Evolva, Switzerland (Eichenberger et al., 2017).

Transporter genes *PUP8* and *PUP1* were expressed from the PGK1 promoter and ADH2 terminator. Coding DNA sequences (CDS) of *PUP8* and *PUP1* were USER fused with PGK1, ADH2 and homologous recombination tags (HRTs) (C and D tags) and cloned into pNB1u plasmid for *in vivo* homologous recombination assembly. USER fusion was done as described previously (Nour-Eldin et al., 2010), using primers shown in Supplementary Table S3. HRTs C and D were obtained by PCR amplification from pEVE2177 plasmid (Eichenberger et al., 2017). For empty vector control, a non-coding DNA sequence (1 kb bp) was used instead of the transporter genes. For the PUP8 localization study, Venus was fused to the C-terminus of PUP8 (PUP8-Venus). *PUP8-Venus* was USER fused together with PGK1, ADH2, C-tag and D-tag into pNB1u plasmid for *in vivo* homologous recombination assembly.

pNB1u vector carrying the different expression cassettes were assembled with pEVE2176 plasmid backbone (carrying *PcUGT88F2* flanked by B and C HRTs) into multi-expression plasmid (with HIS3 as selection marker) using *in vivo* homologous recombination, as described previously (Eichenberger et al., 2017). Helper plasmids carrying fragments required for replication, maintenance and selection, and HRTs required for the assembly, are shown in Supplementary Table S4. Plasmids constructed in this study are also shown in Supplementary Table S4. When UGT and PUP8/PUP1/non-coding sequence is introduced into phloretin pathway strain DBR2, the strain becomes PHZ_PUP8, PHZ_PUP1, or PHZ_control phlorizin-producing strain, respectively.

### 2.5 Media and yeast culture conditions

Unless stated otherwise, synthetic complete drop-out medium without uracil (SD-Ura), without histidine (SD-His) or without uracil and histidine (SD-Ura-His) supplemented with 2 % glucose was used to grow yeast cells for the yeast experiments in this study. SD medium chemicals were purchased from Sigma-Aldrich (St. Louis, Missouri, USA) and medium was prepared according to the manufacturer’s instructions. SD-Ura medium was prepared with 6.7 g/L yeast nitrogen base without amino acids, 1.92 g/L Yeast Synthetic Drop-out Medium Supplement without uracil and 20 g/L glucose. To prepare SD-Ura-His medium, 1.39 g/L Yeast Synthetic Drop-out Medium Supplement without uracil, histidine, leucine and tryptophan was used, and supplemented with 76 mg/L tryptophan and 380 mg/L leucine.

### 2.6 Phlorizin production and sample preparation

For phlorizin production, EnPump200 substrate with enzyme reagent (Reagent A) (EnPresso GmbH, Germany) was used for slow release of glucose to simulate fed-batch conditions. Precultures were inoculated in 3 ml SD-Ura-His medium with 2 % glucose from a single colony in triplicates and incubated at 30 °C with shaking at 250 rpm for 24 h. Main cultures were inoculated in 50 ml SD-Ura-His with 60 g/L EnPump substrate (and 0.1 % Reagent A, glucose-releasing enzyme) as a carbon source in 250 ml shake flask. The initial OD_600_ was 0.1. The flasks were incubated at 30 °C with shaking at 250 rpm for 120 h.

To analyze phlorizin production, 1 ml sample was harvested starting from 48 h after inoculation until 120 h, at an interval of 24 h. The culture OD_600_ was measured and the sample was centrifuged in four replicates and supernatant was collected. Sinigrin glucosinolate was used as internal standard for quantification. The supernatant was filtered through a 0.22 μm filter plate (MSGVN2250, Merck Millipore) and then analyzed for extracellular phlorizin production by LC-MS/MS as described below.

### 2.7 Phlorizin quantification by LC-MS/MS

Oocyte or yeast samples were subjected to analysis by liquid chromatography coupled to mass spectrometry. Chromatography was performed on an Advance UHPLC system (Bruker, Bremen, Germany). Separation was achieved on a Kinetex 1.7u XB-C18 column (100 x 2.1 mm, 1.7 μm, 100 Å, Phenomenex, Torrance, CA, USA). Formic acid (0.05 %) in water and acetonitrile (supplied with 0.05 % formic acid) were employed as mobile phases A and B respectively. The elution profile was: 0-0.1 min, 5 % B; 0.1-1.0 min, 5-45 % B; 1.0-3.0 min 45-100 % B, 3.0-3.5 min 100 % B, 3.5-3.55 min, 100-5 % B and 3.55-4.7 min 5 % B. The mobile phase flow rate was 400 μl min^-1^. The column temperature was maintained at 40 °C. The liquid chromatography was coupled to an EVOQ Elite TripleQuad mass spectrometer (Bruker, Bremen, Germany) equipped with an electrospray ion source (ESI). The instrument parameters were optimized by infusion experiments with pure standards. The ion spray voltage was maintained at +5000 V and −3000 V, in positive and negative ion mode, respectively. Cone temperature was set to 350 °C and cone gas to 20 psi. Heated probe temperature was set to 250 °C and probe gas flow to 50 psi. Nebulizing gas was set to 60 psi and collision gas to 1.6 mTorr. Nitrogen was used as probe and nebulizing gas and argon as collision gas. Active exhaust was constantly on. Multiple reaction monitoring (MRM) was used to monitor analyte molecular ion ➔ fragment ion transitions. Transition for phlorizin was optimized by direct infusion experiments into the MS source. Transitions for Sinigrin and 4MTB were previously reported (Crocoll et al., 2016). Detailed values for mass transitions can be found in Supplementary Table S5. Both Q1 and Q3 quadrupoles were maintained at unit resolution. Bruker MS Workstation software (Version 8.1.2, Bruker, Bremen, Germany) was used for data acquisition and processing. Linearity in ionization efficiencies were verified by analyzing dilution series.

## 3. Results

### 3.1 Identification of phlorizin transporter

An indexed full-length cDNA library of *Arabidopsis* transporters optimized for expression in *Xenopus* oocytes was previously used to identify the first glucosinolates transporters (GTR1 and GTR2) by screening pools consisting of 10 genes per pool for the uptake of glucosinolates in *Xenopus* oocytes (Nour-Eldin et al., 2012). The number of genes in this library was recently increased to 600 genes (unpublished). In this study, we screened this second-generation transporter library for phlorizin transport using LC-MS-based uptake assay in *Xenopus* oocytes (Jørgensen et al., 2017a).

The 600 genes were screened for phlorizin uptake in 75 pools, each containing equal amounts of 8 *in vitro* transcribed cRNAs. A minimum of ten pools (80 genes) was screened at a time in the same batch of oocytes. GTR1-expressing oocytes were included as batch-quality control, and the membrane-impermeable glucosinolate 4MTB (4-methylthiobutyl) was mixed with phlorizin in the uptake assays to control for membrane integrity of oocytes.

Oocytes expressing pool 45 accumulated 122 pmol of phlorizin, whereas oocytes expressing other pools of genes and mock-injected oocytes did not accumulate detectable levels of phlorizin (Figure 1A). Subsequently, pool 45 was split up into two subpools (4 genes in each) and assayed. Phlorizin transport activity was detected only in the second half of pool 45, and this sub-pool was deconvoluted by expressing and assaying each gene individually (Figure 1B and 1C). This identified AT4G18195 as a phlorizin transporter. AT4G18195 is a member of the <u>pu</u>rine <u>p</u>ermease (PUP) transporter family (Gillissen et al., 2000; Jelesko, 2012), and it has been annotated as AtPUP8 (Schwacke et al., 2003).

**Figure 1.**
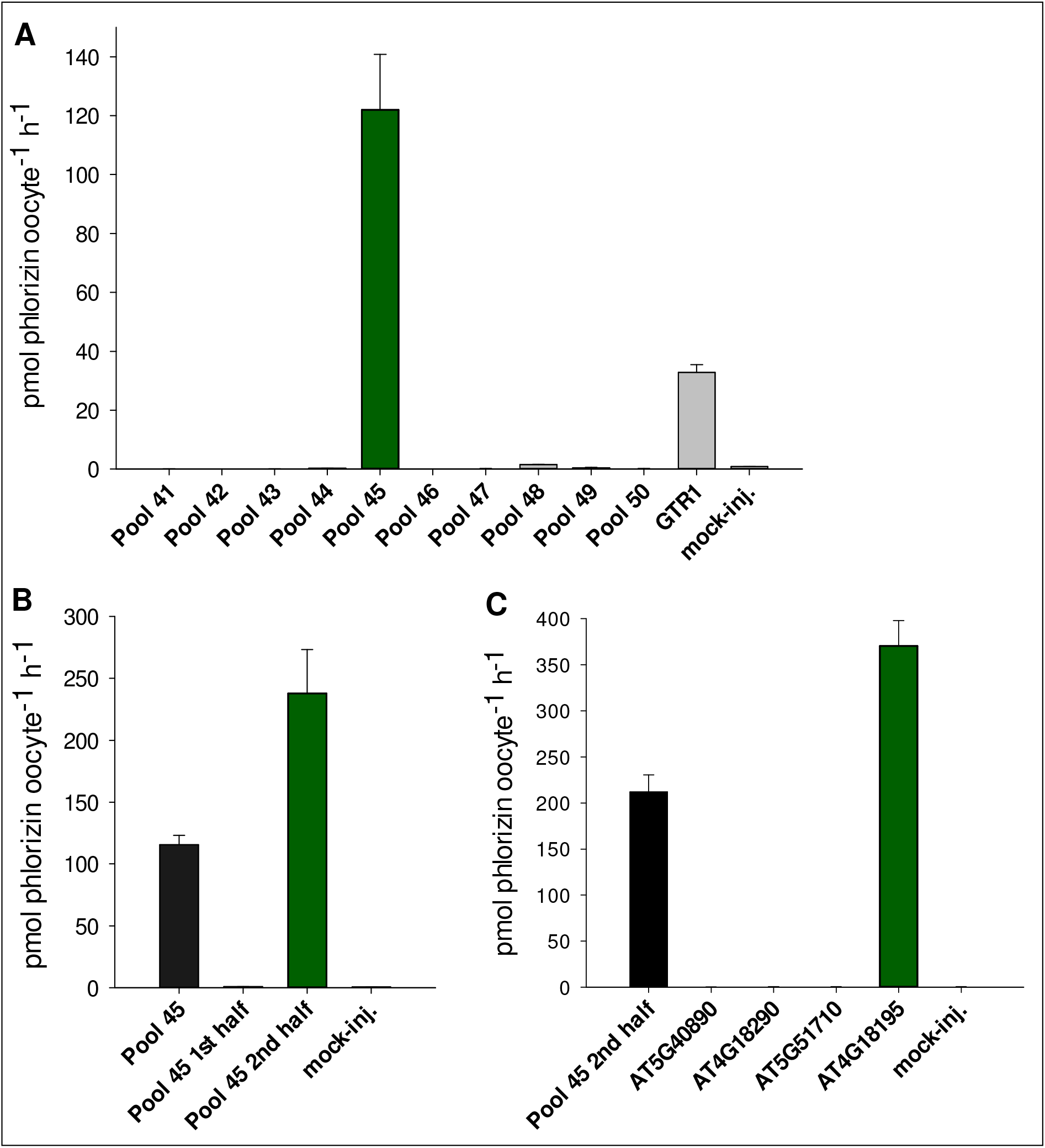
Functional screening of *Arabidopsis* cDNA transporter library for phlorizin transport in *Xenopus* oocytes. A total of 600 genes were screened in pools of 8 genes per pool, 75 pools in total. Only 10 pools (from pool 41 to pool 50) are shown in this figure. (**A**) cRNA of 8 individually *in vitro* transcribed genes was pooled and expressed in 15 oocytes, and transport activity was measured in the presence of 0.5 mM phlorizin (pH 5.0). Phlorizin accumulation within oocytes was quantified using LC-MS/MS analysis. Error bars represent ± s.e. of mean, n = 3 (3 x 5 oocytes). (**B**) The positive phlorizin transporter pool, Pool 45, was split up into two subpools (Pool 45 1^st^ half and Pool 45 2^nd^ half); each subpool contains 4 genes. Subsequently, Pool 45 and the two subpools were expressed in oocytes and transport activity was measured in the presence of 0.5 mM phlorizin. Error bars represent ± s.e. of mean, n = 4 (4 x 5 oocytes). (**C**) Deconvolution of Pool 45 2^nd^ half to identify a phlorizin transporter. The positive subpool and the 4 genes were expressed individually in oocytes, and transport activity was measured in the presence of 0.5 mM phlorizin (pH 5.0). Error bars represent ± s.e. of mean, n = 3 (3 x 5 oocytes).

### 3.2 Biophysical Characterization of PUP8

PUP8 is a 394 amino acid protein with 10 predicted transmembrane spanning domains (Supplementary Figure S1). The PUP family in *Arabidopsis* consists of 21 genes but only PUP1 and PUP2 have been extensively characterized in heterologous systems and shown to transport ring-containing substrates such as the nucleobase adenine and specialized metabolites such as pyridoxine (vitamin B6) and cytokinins (Burkle et al., 2003; Gillissen et al., 2000; Jelesko, 2012; Szydlowski et al., 2013). PUP14 was recently shown to transport cytokinin when expressed in tobacco protoplasts (Zürcher et al., 2016). Additionally, PUP family members from tobacco (NtNUP1), rice (OsPUP7) and coffee (CcPUP1 and CcPUP5) have been shown to transport nicotine, cytokinins and adenine, respectively, when expressed in yeast (Hildreth et al., 2011; Kakegawa et al., 2019; Qi and Xiong, 2013). Thus, the transport activity of PUP8 towards dihydrochalcones ascribes a novel function to the PUP family.

#### 3.2.1. PUP8 is the only transporter in the family that transports phlorizin

Substrate specificity among closely related transporters are typically similar. Accordingly, within the PUP family, AtPUP1 and AtPUP2 can both transport adenine and cytokinins (Burkle et al., 2003; Gillissen et al., 2000). Hence, the identification of PUP8 suggested that other phlorizin transporters may exist in the PUP family. Of the 21 *Arabidopsis* members, 14 were already present in our transporter library. We cloned the remaining 7 PUPs into *Xenopus* oocyte expression vector (pNB1u) and tested the 21 PUPs individually for phlorizin transport activity in *Xenopus* oocytes. Only PUP8 showed phlorizin uptake, whereas the rest of the family, including two closely related homologs PUP7 and PUP21 (which share 76.4 % and 75.8 % amino acid identity with PUP8, respectively), did not show phlorizin uptake (Figure 2A and 2B).

**Figure 2.**
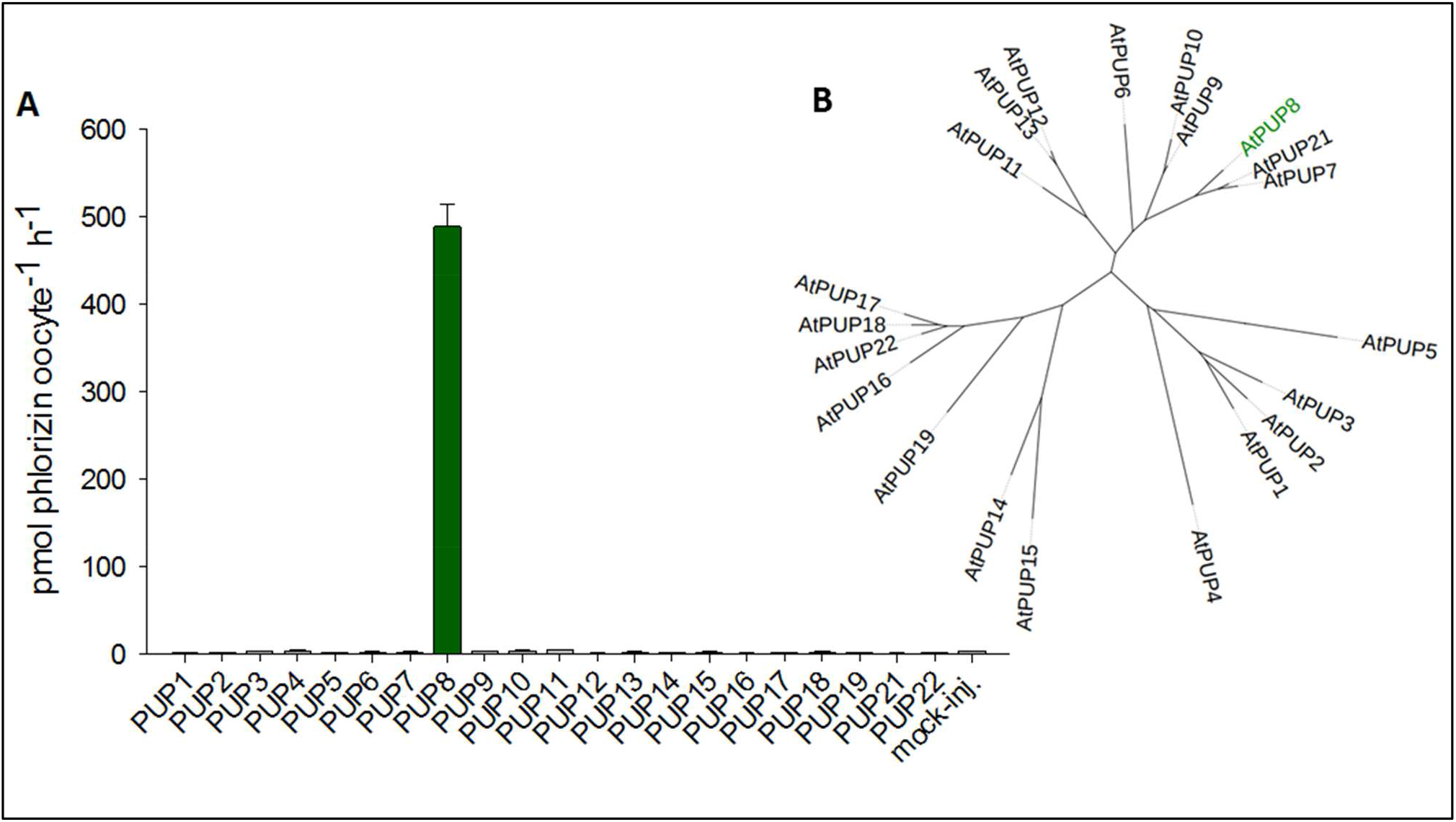
Testing the *Arabidopsis* PUP family transporters for phlorizin uptake in *Xenopus* oocytes. (**A**) 21 AtPUPs were expressed individually in oocytes and transport activity was measured in the presence of 0.5 mM phlorizin (pH 5.0). Phlorizin accumulation within oocytes was quantified using LC-MS/MS analysis. Error bars represent ± s.e. of mean, n = 3 (3 x 3 oocytes). (**B**) Phylogenetic tree of the *Arabidopsis* PUP family. The tree was constructed by the neighbor-joining method. AtPUP8 is shown in green.

To investigate whether the negative results for phlorizin transport were due to lack of expression of other PUP proteins in *Xenopus* oocytes, we tested some of the PUPs (PUP1, 8, 10, 11, 14 and 21) for adenine uptake. PUP1, PUP8, PUP10 and PUP11 transported adenine into oocytes, whereas PUP14 and PUP21 did not transport adenine (Supplementary Figure S2). This result indicates that PUP proteins are generally functional in oocytes and expands the list of verified adenine transporting PUPs from PUP1 and PUP2 to also include PUP8, 10 and 11. However, in the absence of activity for PUP21, we cannot conclude whether lack of phlorizin uptake by the close homologs of PUP8 is due to distinct substrate preference or alternatively non-functional expression. Based on these results, we focus on PUP8 in the remainder of this study.

#### 3.2.2. PUP8 does not transport phlorizin against a concentration gradient

A number of PUP family members have so far been characterized as active secondary transporters that utilize the electrochemical proton gradient to drive transport of their substrates uphill against a concentration gradients (Burkle et al., 2003; Gillissen et al., 2000; Kakegawa et al., 2019; Szydlowski et al., 2013). Active transport is characterized by the transport of a substrate against its concentration gradient. To investigate whether PUP8 transports phlorizin actively, we conducted a 3 h time-course uptake assay in PUP8-expressing oocytes and measured phlorizin accumulation. We found that the transport activity was linear during the first 30 min, and then it saturated when intracellular phlorizin concentration reached the extracellular phlorizin concentration (Figure 3). This shows that PUP8 in oocytes cannot transport phlorizin actively against a concentration gradient and thereby indicates that it functions as a facilitator that allows phlorizin transport via a passive transport mechanism.

**Figure 3.**
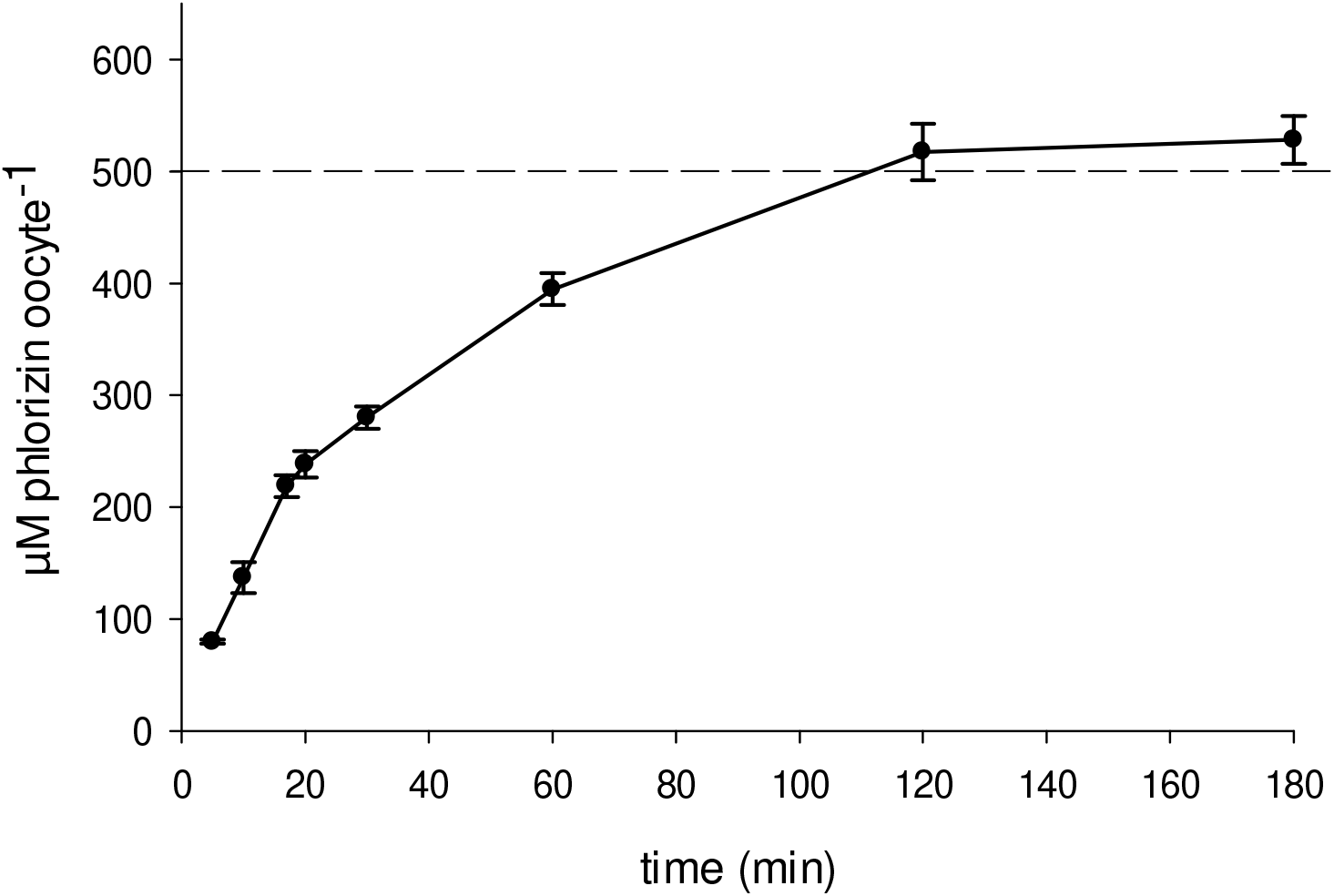
A time-course uptake of phlorizin by PUP8 in *Xenopus* oocytes. PUP8-expressing oocytes were incubated in kulori buffer containing 0.5 mM phlorizin (pH 5.0) for 8 different time points, ranging from 5 min to 180 min. Phlorizin accumulation within oocytes was quantified using LC-MS/MS analysis and plotted against incubation time. Error bars represent ± s.e. of mean, n = 5 (5 x 3 oocytes). The dashed line indicates the phlorizin concentration in the external medium.

#### 3.2.3. Is phlorizin transport by PUP8 dependent on a proton gradient?

A passive transport mechanism implies that PUP8—in contrast to PUP1 (Gillissen et al., 2000; Szydlowski et al., 2013)—transports phlorizin without coupling to proton movement. To investigate this, we tested uptake in oocytes at different extracellular pH (4.5 – 7.5). Phlorizin uptake by PUP8 was not significantly changed when the extracellular pH was increased from 4.5 to 5.5 and 6.5 (Figure 4A). The transport activity was slightly decreased at pH 7.5 by 28 % (compared to pH 5.5). To further investigate the dependency of PUP8 on a proton gradient, a phlorizin uptake was tested in the presence or absence of the proton uncoupler CCCP (carbonyl cyanide m-chlorophenyl-hydrazone). We included a known proton-symporter—GTR1—as a positive control of proton-coupled transport. The addition of CCCP (100 μM) decreased GTR1 transport activity by 84 % (from 713 to 111 pmol 4MTB/oocyte) (Figure 4C), whereas the phlorizin transport by PUP8 was decreased only by 29 % (from 542 to 383 pmol/oocyte) (Figure 4B). Finally, we investigated directly whether PUP8-mediated phlorizin transport is coupled to the movement of protons by subjecting PUP8-expressing oocytes to two-electrode voltage-clamp (TEVC) electrophysiological measurements. Since phlorizin is non-charged, co-transport with even a single proton would induce a negative current in the TEVC measurements. However, phlorizin transport by PUP8 did not induce any negative currents that could correspond with proton-cotransport (Supplementary Figure S3A and B).

**Figure 4.**
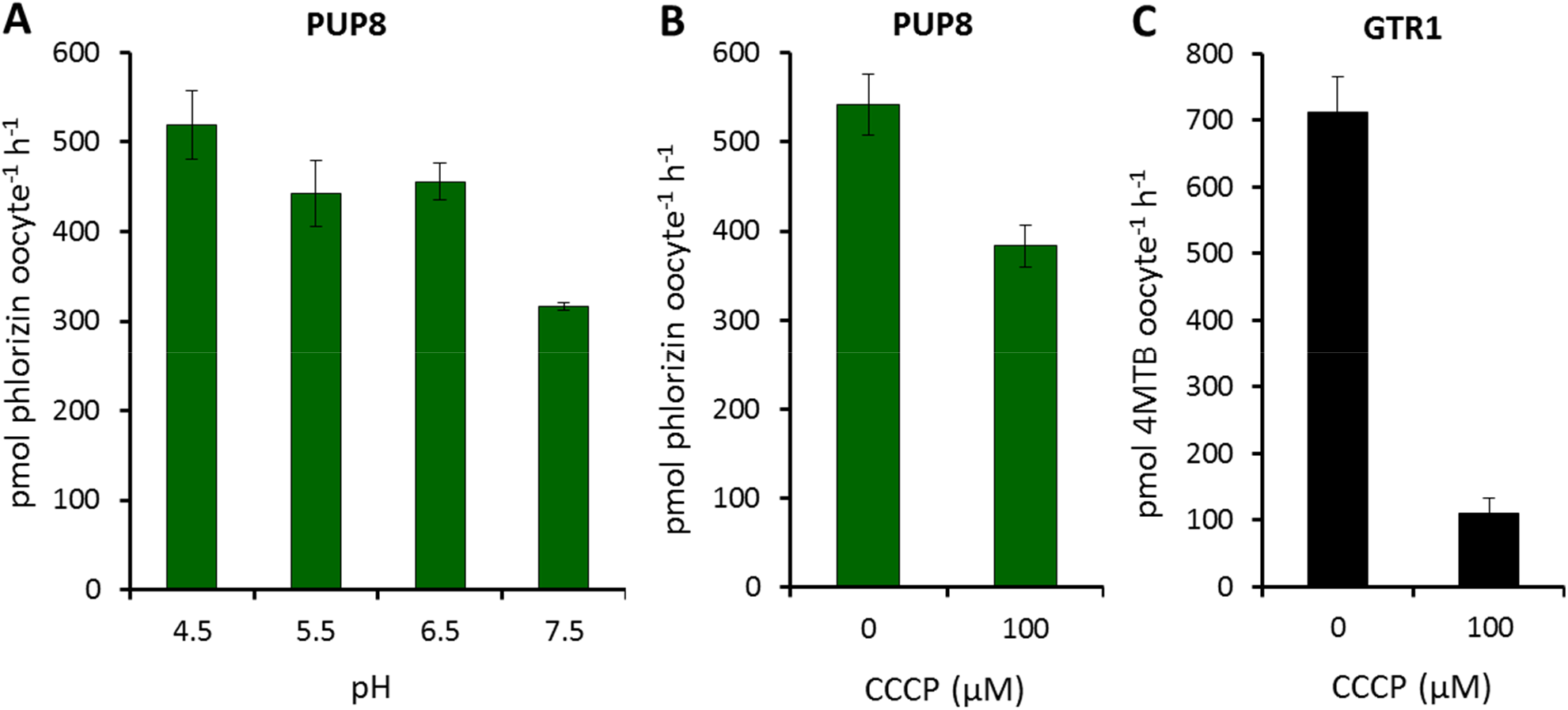
Effect of extracellular proton concentration on PUP8-mediated phlorizin transport in *Xenopus* oocytes. (**A**) Phlorizin transport by PUP8 at different extracellular pH. PUP8-expressing oocytes were incubated in 0.5 mM phlorizin containing buffer at four different pH (4.5, 5.5, 6.5 and 7.5) for 1 h. Phlorizin accumulation within oocytes was quantified using LC-MS/MS analysis. Error bars represent ± s.d. of mean, n = 4 - 5 (4 - 5 x 3 oocytes). Effect of CCCP on PUP8-mediated phlorizin uptake (**B**) or GTR1-mediated 4MTB uptake (**C**). PUP8 and GTR1-expressing oocytes were incubated in 0.5 mM phlorizin and 0.1 mM 4MTB containing kulori buffer (pH 5.0) respectively, in the presence or absence of 0.1 mM CCCP for 1 h. Phlorizin and 4MTB accumulated within the oocytes were quantified using LC-MS/MS analysis. Error bars represent ± s.d. of mean, n = 3 - 4 (3 - 4 x 3 oocytes).

#### 3.2.4. Determining transport kinetics of phlorizin transport of PUP8

Kinetic characterization was performed to estimate the Km of PUP8-mediated phlorizin transport in *Xenopus* oocytes. We chose 15 min assay for kinetic characterization of PUP8-mediated phlorizin uptake at increasing phlorizin concentrations at pH 5. At 15 min PUP8 mediated phlorizin uptake was linear and assumed to represent initial transport rates (Figure 3). The data were fitted to the Michaelis-Menten equation, which estimated a Km value for phlorizin uptake by PUP8 to 296 ± 39 μM (Figure 5). As phlorizin transport by PUP8 was not electrogenic, all our characterizations were performed via LC-MS/MS-based transport assays. Such cumulative transport assays mean that the kinetic characterization is not obtained using true initial transport rates. However, by choosing an assay time within the linear range, we believe that the affinity is estimated to the best of the current technical capability.

**Figure 5.**
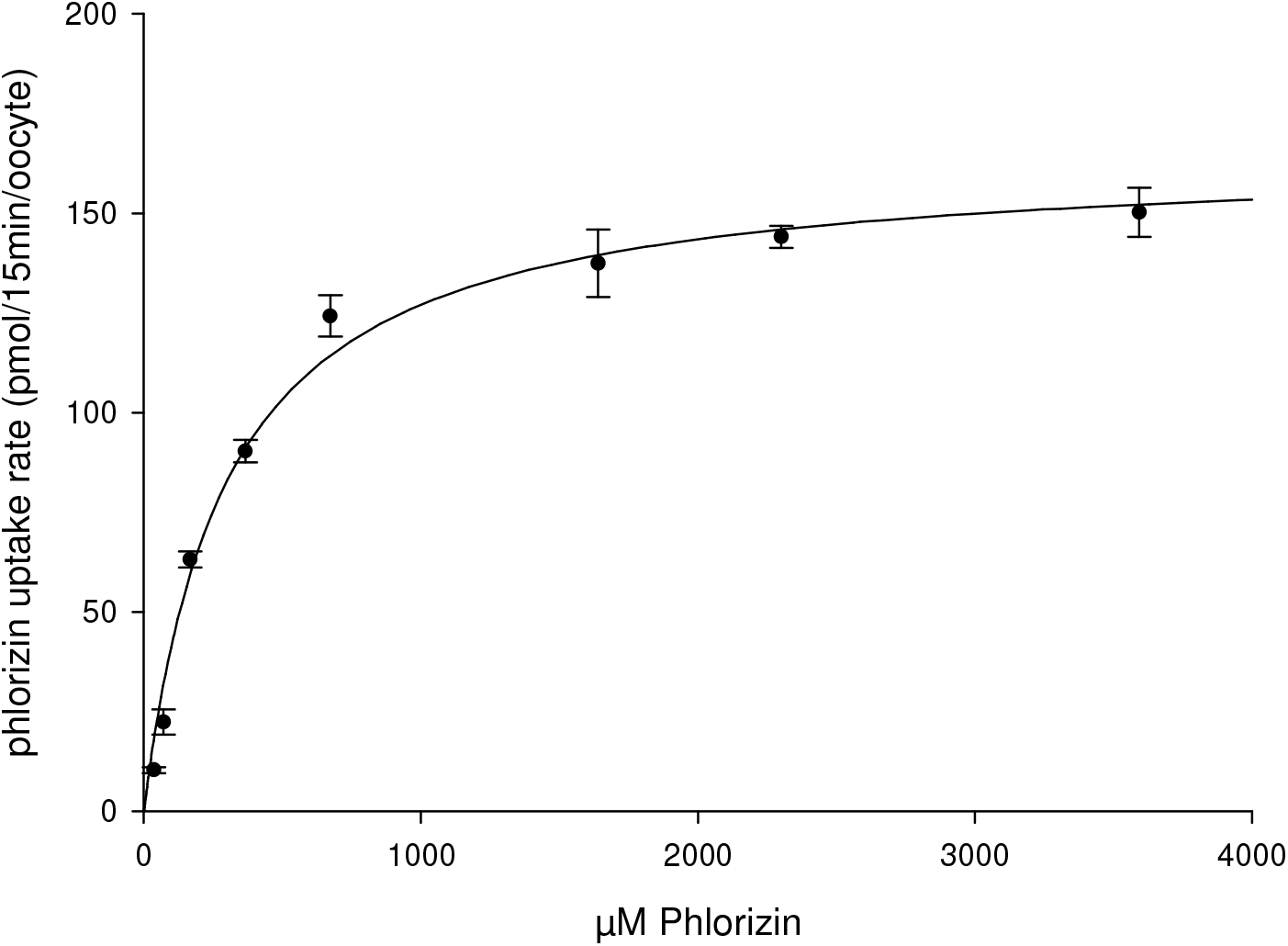
Kinetic characterization of PUP8-mediated phlorizin transport. PUP8-expressing oocytes were assayed at 8 different phlorizin concentrations for 15 min at pH 5.0. The data were fitted to the Michaelis-Menten equation. Phlorizin accumulation within oocytes was quantified using LC-MS/MS analysis. Error bars represent ± s.e. of mean, n = 4 (4 x 3 oocytes).

#### 3.2.5. Directionality of phlorizin transport of PUP8

The apparent passive transport mechanism of PUP8 suggested that it is a facilitator that could accommodate bidirectional phlorizin transport. To test whether PUP8 can also export phlorizin, we performed phlorizin injection-based export assay in *Xenopus* oocytes. We measured phlorizin efflux from phlorizin-injected oocytes over time, by quantifying both intracellular and extracellular phlorizin content. The injected intracellular phlorizin amount decreased significantly over time in PUP8-expressing oocytes, and simultaneously the amount of phlorizin in the extracellular solution increased (Figure 6). 14 h after phlorizin injection, PUP8-expressing oocytes had exported 65 % of the injected phlorizin. By 22 h, 75 % of the injected phlorizin had been exported (Figure 6). In contrast, mock-injected oocytes did not display significant phlorizin efflux. To rule out that the export was an artifact induced by expression of a heterologous transporter, PUP1 (which functions in oocytes as an adenine transporter, Supplementary Figure S2) was tested and showed a pattern similar to the mock-injected oocytes (Figure 6). Furthermore, we verified that phlorizin export in PUP8-expressing oocytes was specific to phlorizin as 4MTB was not exported from PUP8-expressing oocytes (Supplementary Figure S4). These results indicate that PUP8 is capable of mediating both phlorizin import and export. The direction of net transport likely depends on the phlorizin concentration gradient, which is consistent with a uniport transport mechanism.

**Figure 6.**
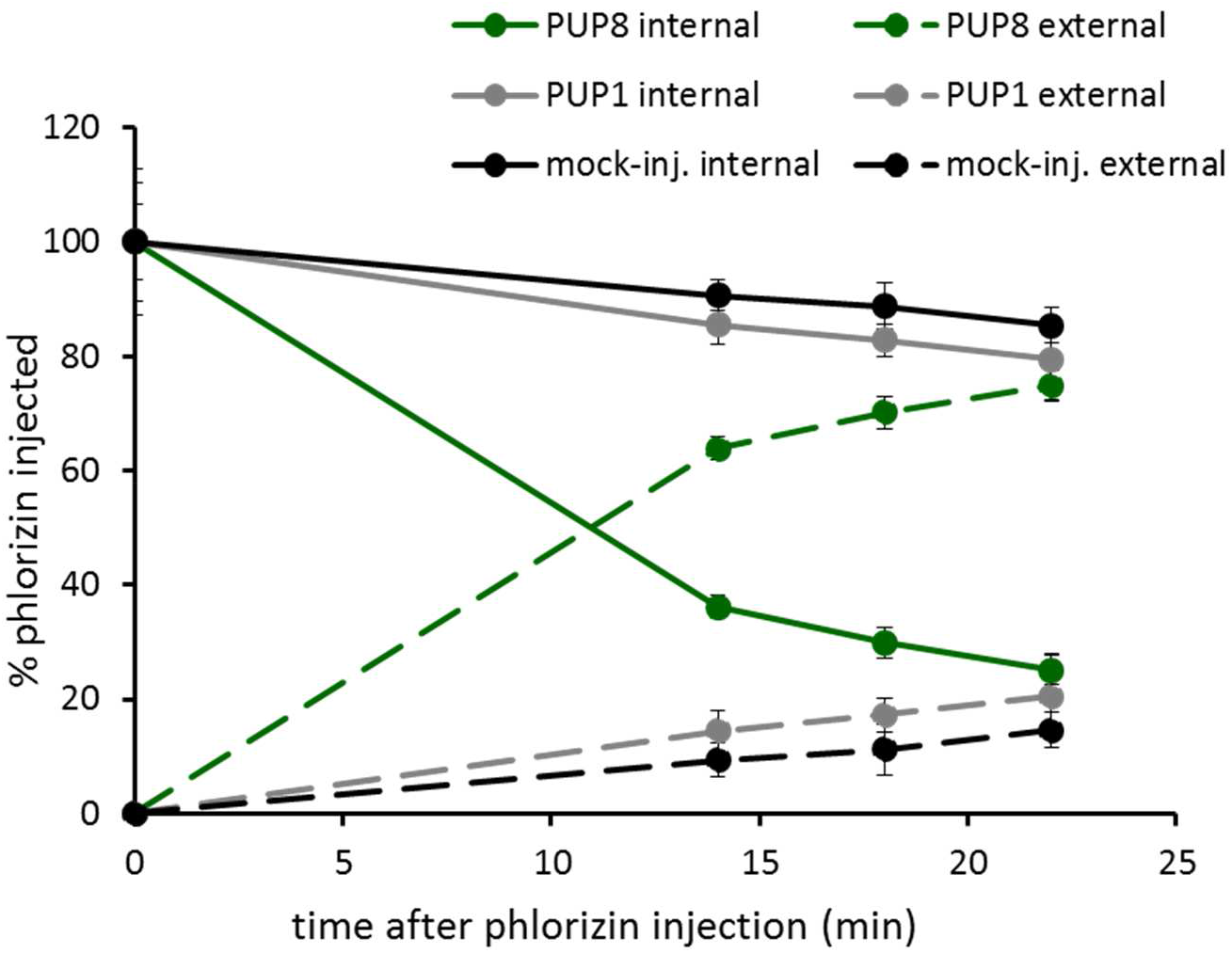
Injection-based export assay of phlorizin by PUP8 in *Xenopus* oocytes. Oocytes expressing PUP8, PUP1, or mock-injected oocytes were injected with phlorizin (to obtain initial internal concentration ~0.5 mM) and incubated in a kulori buffer (pH 7.4, without phlorizin). Export transport activity was measured by quantifying intracellular and extracellular phlorizin content, after 14, 18 and 22 h of incubation, using LC-MS/MS analysis. Error bars represent ± s.d. of mean of % phlorizin injected, n = 5 (5 x 3 oocytes).

### 3.3 Subcellular localization and functional expression of PUP8 in yeast

PUP8 was identified and hitherto characterized in *Xenopus* oocytes. A number of PUP family members have previously been functionally characterized in yeast (Burkle et al., 2003; Gillissen et al., 2000; Kakegawa et al., 2019; Qi and Xiong, 2013), suggesting that they are correctly localized in the plasma membrane. However, a subcellular localization study of the tobacco PUP family nicotine transporter NtNUP1 shows that it is not only localized to the plasma membrane, but also to endomembranes (Kato et al., 2015). Thus, before we proceeded to express PUP8 in the phlorizin-producing yeast strain, we sought to verify the subcellular localization of PUP8 in *S. cerevisiae*.

We first studied the expression and localization of PUP8 in yeast by translationally fusing Venus to the C-terminus of PUP8 (PUP8-Venus) and analyzing by confocal microscopy. PUP8-Venus localized predominantly to the plasma membrane and endomembrane, likely the tonoplast (Figure 7A). Next, we tested the functionality of PUP8 in yeast; we performed phlorizin uptake assay by incubating yeast cells expressing either PUP8 or empty vector (control) in a buffer containing 0.5 mM phlorizin for 30 min, and phlorizin accumulation in yeast cells was quantified using LC-MS/MS analysis. PUP8-expressing yeast cells accumulated a considerable amount of phlorizin (644 pmol/O.D/ml), whereas the control cells accumulated low background levels (Figure 7C). These results demonstrated that PUP8 is functionally expressed in yeast’s plasma membrane. However, due to the signals at what appears to be the tonoplast (Figure 7A), we cannot exclude that PUP8 may also facilitate transport of phlorizin across intracellular endomembranes.

**Figure 7.**
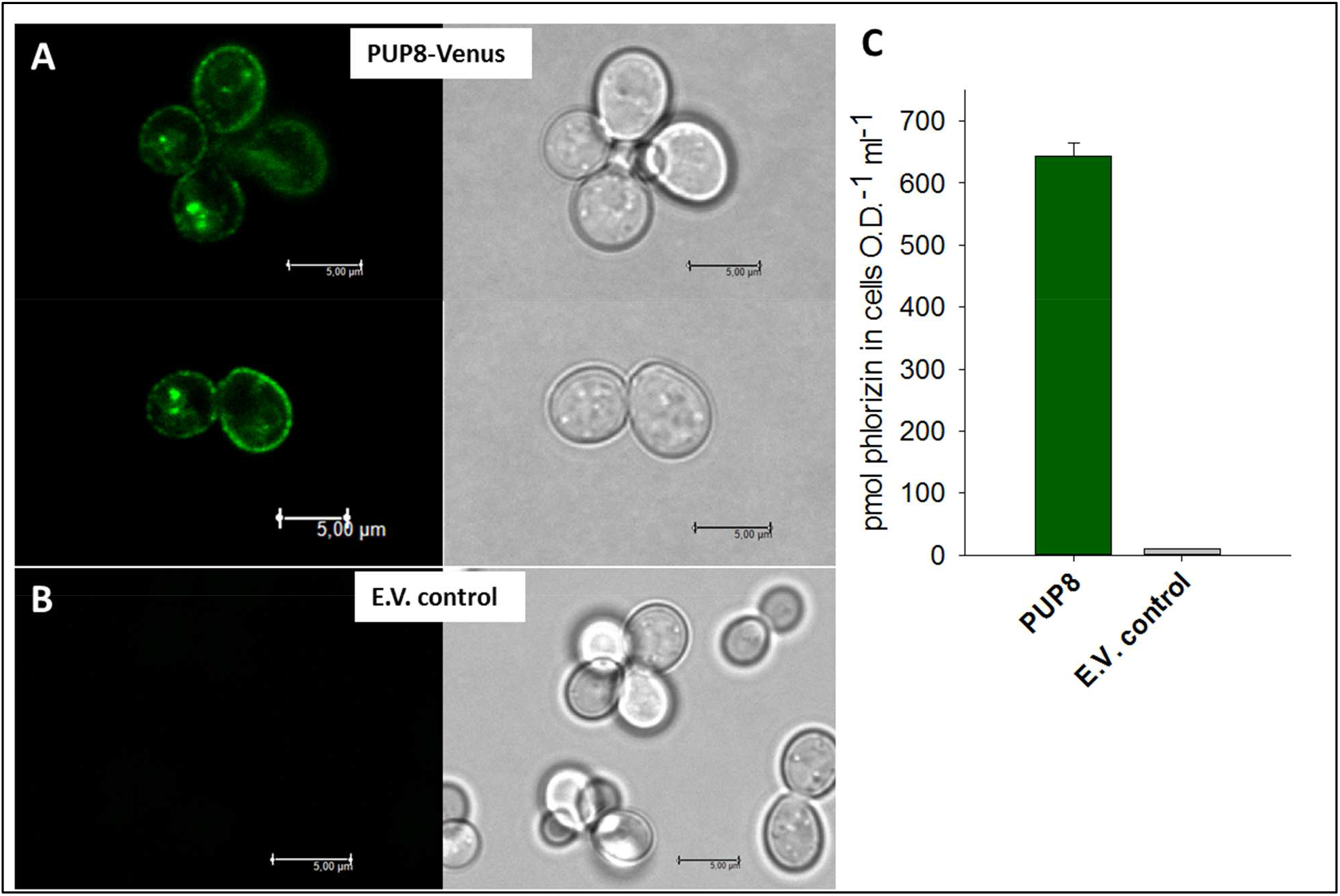
Subcellular localization of PUP8 and its phlorizin transport in *S. cerevisiae*. PUP8-Venus fusion protein was expressed in yeast under the control of the *S. cerevisiae* PGK1 promoter **(A)**. Empty vector control without Venus expressions was used as a negative control **(B).** After overnight growth in SD-His medium (2 % glucose), cells were analyzed by confocal microscopy (Leica SP5-X). The left image was taken under Venus excitation light; the corresponding bright-field image is presented on the right side. All the scale bars represent 5 μm. **(C)** Phlorizin uptake assay in *S. cerevisiae*. Yeast cells expressing PUP8 or empty vector (E.V.) control were incubated in a phosphate buffer (0.1 M HK2PO4, pH 5.0) containing 0.5 mM phlorizin for 30 min. Following washing the pellet twice, phlorizin accumulation within the yeast cells was quantified using LC-MS/MS analysis. Error bars represent ± s.d. of mean, n = 4 (4 x 3 transformants).

### 3.4 Expression of PUP8 increased phlorizin production in S. cerevisiae

The phlorizin-producing strain PHZ3, described by Eichenberger et al. (2017), comprises a set of two HRT plasmids, the first of which carries the phloretin pathway consisting of six genes (as in DBR2), whereas the UGT gene (PcUGT88F2) is expressed from a second plasmid. To investigate whether the expression of PUP8 increases phlorizin production, we expressed PUP8 and PcUGT88F2 on the second HRT plasmid in strain DBR2, making strain PHZ_PUP8. For negative control, the CDS of PUP8 was replaced by a non-coding sequence and then expressed in strain DBR2, making strain PHZ_control. The strains were cultured in a simulated fed-batch medium in shake flasks and the extracellular phlorizin production was quantified by LC-MS/MS.

The PHZ_PUP8 strain produced significantly higher extracellular levels of phlorizin than the negative control strain PHZ_control at all time points (Figure 8A). After 96 h of fermentation, the PHZ_PUP8 strain produced 75 ± 4 mg/L extracellular phlorizin, which is 83 % higher than the control strain (41 mg/L). To exclude the possibility that the increased phlorizin titer in PHZ_PUP8 strain was due to improved adenine import or an artifact induced by the expression of purine permease protein rather than facilitated phlorizin export, we expressed the adenine transporter PUP1 (PHZ_PUP1) and analyzed phlorizin production. Expressing PUP1 did not improve the phlorizin production (Figure 8A). From the time-dependent phlorizin production assay, the optimum production appeared to be at 96 h. To account for potential effects on growth, we normalized the phlorizin titer to the optical density. At 96 h, expression of PUP8 resulted in a 79 % increase in phlorizin specific yield (mg/L/OD) as compared to the control strains (Figure 8B). We also showed that external application of phloretin or phlorizin, at concentrations at least 3-fold higher than the production level of the strains, did not affect the yeast growth (data not shown).

**Figure 8:**
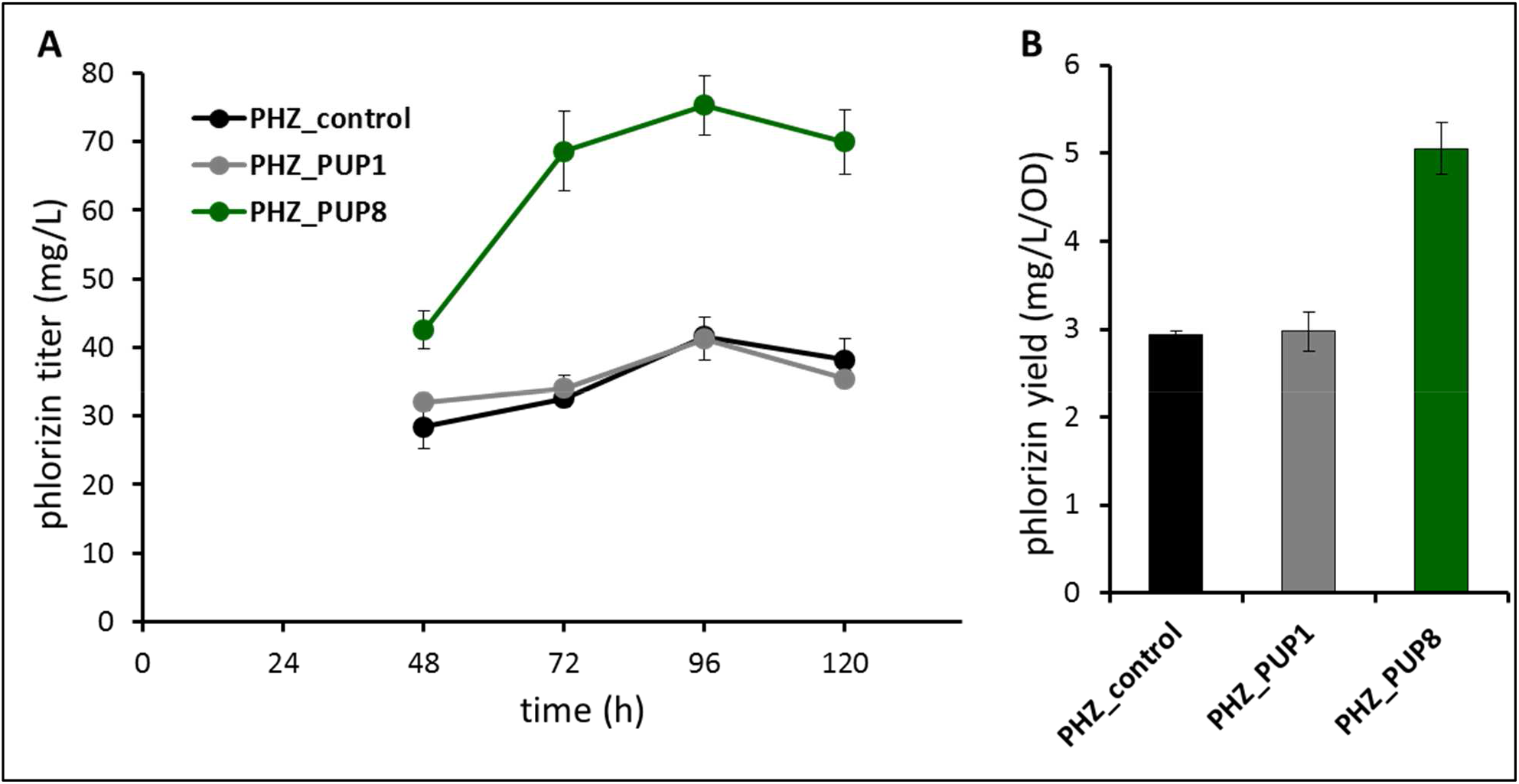
Effect of PUP8 expression on phlorizin production in *S. cerevisiae*. Phlorizin-producing yeast strains expressing PUP8 (PHZ_PUP8), PUP1 (PHZ_PUP1) or empty vector control (PHZ_control) were cultivated in SD-Ura-His medium with 60 g/L EnPump substrate (with 0.1 % enzyme reagent) for slow glucose release to simulate fed-batch conditions in shake flasks. Subsequently, samples were harvested at 48, 72, 96 and 120 h after inoculation, and external phlorizin production was quantified using LC-MS/MS analysis. Time-dependent phlorizin titer (mg/l) **(A)**, and phlorizin specific yield at 96 h after inoculation **(B)** are shown. Error bars represent ± s.d. of mean, n = 4.

## Discussion

Transport-engineering represents an emerging technology for increasing production in bioengineering through improving substrate supply or by facilitating efflux of the final product into the growth medium (Boyarskiy and Tullman-Ercek, 2015; Nour-Eldin and Halkier, 2013). To realize the potential of transport-engineering, we addressed the need to exploit the vast repository of plant transporters of specialized metabolites and to study their effects on yield through case studies (Borodina, 2019; Larsen et al., 2017b). As a first step, we establish unbiased brute-force screening of plant transporter libraries in *Xenopus* oocytes as a viable approach for identifying plant transporters for bioengineering purposes. Hence, it is timely to discuss how the experiences made in this study will shape future screening endeavors.

### Library design – continued screening of *Arabidopsis* library and in parallel build targeted hotspot library

A pertinent question is whether to invest time and labor into building new transportome-wide species-specific transporter libraries for future identification of transporters of novel compounds. The large ABC transporter family has long been known to encode detoxifying transporters capable of exporting a wide range of xenobiotics (Kang et al., 2011) and recently, members from the NPF was shown to transport specialized plant metabolites with highly varying chemical structures (Jørgensen et al., 2017b; Larsen et al., 2017a; Nour-Eldin et al., 2012). Similarly, the substrate spectrum of the PUP family was recently expanded from purine nucleobase substrates (Gillissen et al., 2000) to include plant specialized metabolites such as pyridoxine, nicotine, benzylisoquinoline alkaloids and now phlorizin (Dastmalchi et al., 2019; Jelesko, 2012; Kato et al., 2015; Szydlowski et al., 2013). The large disparity in chemical structure between these substrates indicates that the substrate spectrum of plant transporter families may be very large. Accordingly, we find it worthwhile to screen the already-established *Arabidopsis* library (600 transporter genes) for activity towards future target compounds. As shown in this study, rapid screening of this library may identify a transporter that can serve the immediate needs of ongoing bioengineering projects and holds potential for identifying additional transporter families with substrate specificity towards specialized metabolites. In parallel, smaller dedicated libraries should be built from plants producing the target compounds. These libraries should consist of hotspot families known to encode transporters of specialized plant metabolites and that express well in *Xenopus* oocytes (such as the NPF and the PUP families). We believe that these transport libraries will form a strong foundation for future transporter identification for synthetic biology purposes.

### Assay design – using import assays for identifying exporters

Exporters, crucial for efficient secretion of final products, are particularly challenging to screen for directly in any model system. *Xenopus* oocytes provide a unique advantage as target substrates can be delivered to the cytosol via injection. For high-throughput screening of large transporter libraries, the need for an extra injection is labor intensive and imposes a sustained—hard-to-fulfill—requirement for oocytes of high robustness. Additionally, when characterizing the directionality of PUP8, we detected PUP8 mediated phlorizin export only after 14 h past compound injection (Figure 6). In comparison, some compounds such as sucrose export by SWEET transporters was detected already after 5 minutes of substrate injection (Chen et al., 2010, 2012; Lin et al., 2014). This difference in assay-time before export could be detected indicates that the time required for intracellular movement of injected substrates to the membrane for transporter-mediated export may vary in a substrate-dependent manner. After 48 h, PUP8-expressing and mock-injected oocytes had both exported almost all injected phlorizin (data not shown). Thus, long-term endpoint assays may be unsuitable due to potential endogenous export activity. This unpredictability in assay-time imposes the need to monitor export using laborious time-course assays, which further complicates library screening for exporters using injection-based export assay.

Instead, we identified PUP8 via a straightforward uptake-based screen and then demonstrated bidirectionality by injecting phlorizin in PUP8-expressing oocytes and detecting PUP8-dependent extracellular phlorizin accumulation over time. This aligns well with the elegant uptake-assay used to identify the long-sought SWEET sucrose exporters (Chen et al., 2010). Similarly, the characterization of the H^+^-coupled ZmSUT1 sucrose transporter showed that the sucrose-coupled proton current was reversible and depended on the direction of the sucrose and pH gradient as well as the membrane potential across the transporter (Carpaneto et al., 2005). However, we do not expect that uptake-based assays are suitable for all types of transporters. For example, eukaryotic ABC transporters have long been accepted to act exclusively as exporters (Kang et al., 2011). Therefore, for ABC transporters it may be necessary to build dedicated sub-libraries that can be screened via the injection-based time-course export assays. Thus, when searching among secondary transporters (carriers) for export activity it is possible to use straightforward uptake assays and then investigate whether they possess bidirectionality.

### *Xenopus* oocytes as expression host – best in the class but not necessarily perfect?

The fact that we could express and detect adenine uptake for four different PUP members (Supplementary Figure S2) indicated that *Xenopus* oocytes are generally able to express, fold and target PUP members functionally to the plasma membrane. However, it was expected that PUP8’s close homologs (PUP7 and PUP21, 76.4 % and 75.8 % amino acid identity, respectively) would also exert detectable phlorizin transport. However, despite their homology to PUP8, PUP7 and PUP21 remain as orphan transporters with no experimentally verified substrate. On the other hand, PUP8 appears to possess dual substrate specificity towards phlorizin and adenine (Figure 1C and Supplementary Figure S2). In this context, it is noteworthy that PUP14 was previously shown to mediate cytokinin (trans-zeatin) uptake when expressed in protoplasts from *Nicotiana benthamiana* (Zürcher et al., 2016). Here, PUP14 injected oocytes did not transport adenine (Supplementary Figure S2), which is otherwise a substrate for cytokinin-transporting PUPs (Burkle et al., 2003; Gillissen et al., 2000). Nor could we detect cytokinin (trans-zeatin) uptake for PUP14 when expressed in *Xenopus* oocytes (data not shown). This non-functional expression of PUP members with known (or expected) substrates could be due to a number of reasons, for example, lack of co-factor(s) or interacting partners or that the transporters require activation through posttranslational modification in the *Xenopus* oocyte system. Despite *Xenopus* oocytes representing an extremely well-established system for expressing and characterizing transport proteins, there is room for further optimization. For example, obtaining functional expression of PUP14 in *Xenopus* oocytes represents an interesting case study that may reveal important alterations that may find generic applicability when expressing other PUP members.

### Increasing phlorizin production in yeast through transport-engineering

As one of the prerequisites for cost-efficient heterologous production of valuable small molecules, a microbial chassis is required to export final products to the external growth medium. This enables facile product recovery without the need for costly purification from cell homogenate. Moreover, efficient export can alleviate potential feedback inhibition and auto-toxicity, which otherwise may hamper yield. An open question is at which point in a bioengineering endeavor it would be beneficial to introduce a transporter to improve secretion if at all.

Phlorizin is a hydrophilic compound that requires transport proteins to traverse membrane bilayers. By collecting external growth medium, we showed that the phlorizin-producing yeast strain used in this study secretes phlorizin through endogenous transporters (Figure 8). Previous studies have shown that production levels of target compounds can be increased by improving endogenous transport activities (Fisher et al., 2014; Foo and Leong, 2013). Accordingly, introducing a heterologous plant transporter represents another means to improve export of microbial cell factories with potential for increased yield. Indeed, PUP8 expression significantly increased phlorizin production as compared to the control strains (Figure 8A). However, given the dual substrate specificity of PUP8 (adenine and phlorizin) (Figure 1C and Supplementary Figure S2), we asked whether this increase could be an indirect effect of increased adenine transport activity. Adenine supply is a principal component of yeast extract media that affects recombinant protein production (Zhang et al., 2003). As a control, we show that PUP1 transports adenine but not phlorizin in both *Xenopus* oocytes and yeast. The inability of PUP1 to increase phlorizin production in the phlorizin producing yeast strain (Figure 8) indicates that PUP8-mediated phlorizin export is the direct cause for increased phlorizin production. As a case study on implementing plant transporters in bioengineering, this positive outcome prompted us to consider the underlying causes for this yield increase.

### Phlorizin producing yeast strain may be subject to feedback inhibition

Phlorizin is a known competitive inhibitor of mammalian sodium-coupled SGLT glucose transporters and its aglycone phloretin inhibits the mammalian GLUT transporters (Ehrenkranz et al., 2005). However, the 17 HXT glucose transporters in *Saccharomyces cerevisiae* are neither inhibited by phlorizin nor phloretin (Kasahara et al., 2009). Thus, it is unlikely that the yield increase is correlated with any modulation of sugar transport activity in yeast.

Phlorizin appears to have few cytotoxic effects whereas the penultimate product, the aglycone phloretin is an uncoupler and inhibitor of mitochondrial oxidative phosphorylation (De Jonge et al., 1983). Introduction of the UGT from pear (PcUGT88F2) into the phloretin producing strain reduced phloretin levels by a factor of ~5, but not all phloretin is converted to phlorizin by the UGT (Eichenberger et al., 2017). Thus, it is possible that the remaining phloretin could hamper cell growth. Via a growth curve experiment, we showed that extracellular application of phloretin or phlorizin (0.5 mM) did not affect the growth of yeast cells (data not shown). This indicates that the increased in phlorizin production by PUP8 is not through alleviating phloretin or phlorizin mediated toxicity. This conclusion is supported by the normalized increase in phlorizin production to OD showing little effect on yeast growth.

Increased phlorizin production could rather be due to alleviated feedback inhibition of biosynthetic enzyme(s). It is noteworthy that the enzymatic activities of general phenylpropanoid and flavonoid biosynthesis pathways are regulated by pathway intermediates (Yin et al., 2012). In particular, the enzymatic activity of phenylalanine ammonia-lyase (PAL), the first committed step in the phlorizin biosynthesis, appears sensitive to even very low levels of non-glycosylated flavonols, which are believed to transmit repression by direct inhibition or by inducing secondary suppressive modifications (Yin et al., 2012). Early phlorizin pathway intermediates, such as cinnamic acid and p-Coumaric acid, have also been reported to feedback control the PAL activity (Blount et al., 2000; Lam et al., 2008; Sarma and Sharma, 1999). Moreover, phlorizin analogs (canagliflozin and dapagliflozin) have been reported to be potent inhibitors of human UGTs (Pattanawongsa et al., 2015). To our knowledge, phloretin and phlorizin mediated feedback inhibition of biosynthetic enzymes have not yet been demonstrated. But, based on the observations presented above it is appears that PUP8 mediated export of phlorizin into the growth medium is key for alleviating feedback inhibition leading to increased enzymatic activity.

Thus, in this case study we show the introduction of heterologous bidirectional phlorizin transport activity significantly improved production from a non-optimized pathway in a microbial chassis that secretes the target compound by endogenous transporters. In the context of bioengineering, it may thus be worthwhile to search for transporters of final products early on in parallel with establishing the biosynthesis pathway.

Moreover, we believe that the findings presented in this study raise a number of interesting questions. For example, we claim that introducing heterologous transporters in most transport-engineering papers have focused on improving ABC transporters, phlorizin transporters with different transport mechanisms would provide insights into which transport properties are desirable. It will be interesting to study whether a synergistic improvement in yield would be observed if PUP8 was introduced in a yeast strain where biosynthesis pathway has been optimized.

In conclusion, we screened a large *Arabidopsis* full-length cDNA library in *Xenopus* oocytes and identified and characterized the plant transporter PUP8, capable of transporting the anti-diabetic compound phlorizin across plasma membranes in a heterologous host. PUP8 was the only transporter in this family that could transport phlorizin in *Xenopus* oocytes. PUP8 transported phlorizin bidirectionally, depending on the concentration gradient in *Xenopus* oocytes. We show that PUP8 is a medium-affinity phlorizin uniporter that can transport phlorizin independent of a pH-gradient. PUP8 expression improved phlorizin titer by 83 %, most likely through facilitating product secretion and thereby accelerating the forward reaction of the pathway by creating a sink for the product. The identification of the *Arabidopsis* transporter PUP8 for the exogenous metabolite phlorizin through cDNA library screening shows that such transporter libraries can be screened to identify transporters for other metabolites of interest. This may help overcome the challenge of identifying transporters for the purpose of future transport-engineering in synthetic biology. In the context of providing transporters for transport-engineering purposes, this study shows that plants encode a treasure cove of transporters with unexpected substrate specificities that can be mined efficiently for activity towards high-value natural products.

## Abbreviations

UGT: UDP-dependent-glycosyltransferase;
HRT: homologous recombination tag;
OD_600_: optical density at 600 nm;
LC-MS: liquid chromatography mass spectrometry

## Acknowledgments

We acknowledge the financial support from the Innovation Fund Denmark (grant 76-2014-3 to ZMB) and the Danish National Research Foundation (grant DNRF99 to CC and HN). IM-H and IB acknowledge the financial support from the Novo Nordisk Foundation (Grant Agreement NNF10CC1016517) and from the European Research Council under the European Union’s Horizon 2020 Research and Innovation Programme (YEAST-TRANS Project, Grant Agreement 757384). The authors would like Louise Svenningsen for technical assistance in the laboratory, Carole Duchêne for assisting in subcloning some PUP genes and Michael Hansen for assisting in confocal microscope imaging. We would also like to thank Evolva SA for making a yeast strain and phlorizin pathway plasmids available for this work.

