## Supplementary methods, figures and tables for "Identification and characterization of phlorizin transporter from *Arabidopsis thaliana* and its application for phlorizin production in *Saccharomyces cerevisiae*"

*1.1 Adenine radiotracer uptake assay in X. laevis oocytes*

Radiotracer uptake assay in oocytes was performed as described previously (Nour-Eldin et al., 2006), with some modifications. The assay was performed by incubating expressing oocytes in 1 µCi/ml $^3$H[adenine] (37.6 Ci/mmol, PerkinElmer) containing kulori buffer (pH 5.0) for 10 min. After 10 min of incubation, oocytes were washed 3 times in kulori buffer. Subsequently, oocytes were transferred to scintillation vials (single oocyte/vial) and lysed in 100 µl 10 % (w/v) SDS by vigorous vortexing. 2.5 ml of EcoScint$^{TM}$ scintillation fluid (National Diagnostics) was subsequently added to each vial followed by vortexing. Radioactivity was quantified using liquid scintillation counting (Nour-Eldin et al., 2006).

*1.2 Phlorizin transport assay in S. cerevisiae*

For phlorizin uptake assay in yeast cells, yeast cells were grown in 10 ml SD-His medium with 2 % glucose at 30 °C with shaking at 250 rpm for 16 h. The cells were harvested to an $OD_{600}$ of 20 by centrifugation, washed once and resuspended in 1 ml ice-cold phosphate buffer (0.1 M $HK_2PO_4$ pH 5.0). The assay was started by resuspending the cells in 0.3 ml 0.5 mM phlorizin containing phosphate buffer (pH 5.0) and incubating at 30 °C with shaking at 400 rpm for 30 min. Subsequently, the cells were collected by centrifugation and washed 3 times with 1 ml ice-cold phosphate buffer (pH 5.0). The cells were lysed with 80 % methanol (containing 1.25 µM sinigrin as internal standard) heating at 45 °C with shaking at 1000 rpm for 15 min. The supernatant was collected by centrifugation, diluted in water and filtered through a 0.22 µm filter plate (MSGVN2250, Merck Millipore) and then the filtrate was analyzed by LC-MS/MS.

*1.3 Confocal microscopy*

*S. cerevisiae* yeast cells expressing PUP8-Venus were grown in 3 ml SD-His medium at 30 °C 200 rpm for 15 h. Subsequently, cells were prepared for imaging by mixing with agarose (0.4 % final concentration). The cells were imaged using a confocal microscope (Leica SP5-X) with a 63×/1.2 water immersion objective.

*1.4 Two-electrode voltage-clamp electrophysiology*

Phlorizin or 4MTB induced currents in expressing oocytes were recorded using the two-electrode voltage-clamp technique on automated Roboocyte2 (Multichannel Systems, Reutlingen, Germany). Electrodes were backfilled with a mixture of 3 M KCl and 1.5 M KAcetate. Oocytes were continuously perfused with MES-based ekulori buffer (90 mM NaCl, 1 mM KCl, 1 mM $MgCl_2$, 1 mM CaCl2, 2 mM LaCl3, 5 mM MES pH 5.0). The holding potential was -60 mV. Currents were recorded while clamping the membrane potential of oocytes stepwise from -120 mV to +20 mV in 20 mV increments before and after addition of substrate (2 mM phlorizin or 0.5 mM 4MTB) in MES-based ekulori buffer (pH 5). Electrodes had resistance of 280 - 1000 kΩ. The current-voltage (*IV*) curves were plotted by subtracting currents before substrate addition from currents recorded after substrate addition.

*1.5 Phylogeny analysis*

Phylogeny of the *Arabidopsis* PUP family proteins were analyzed using CLC Genomics Workbench 7.6 (Qiagen). Sequences of the 21 AtPUP genes were obtained from TAIR (https://www.*Arabidopsis*.org) and sequence alignment of proteins was performed using full-length amino acid sequences. The Phylogenetic tree of the AtPUPs was constructed by the neighbor-joining method.

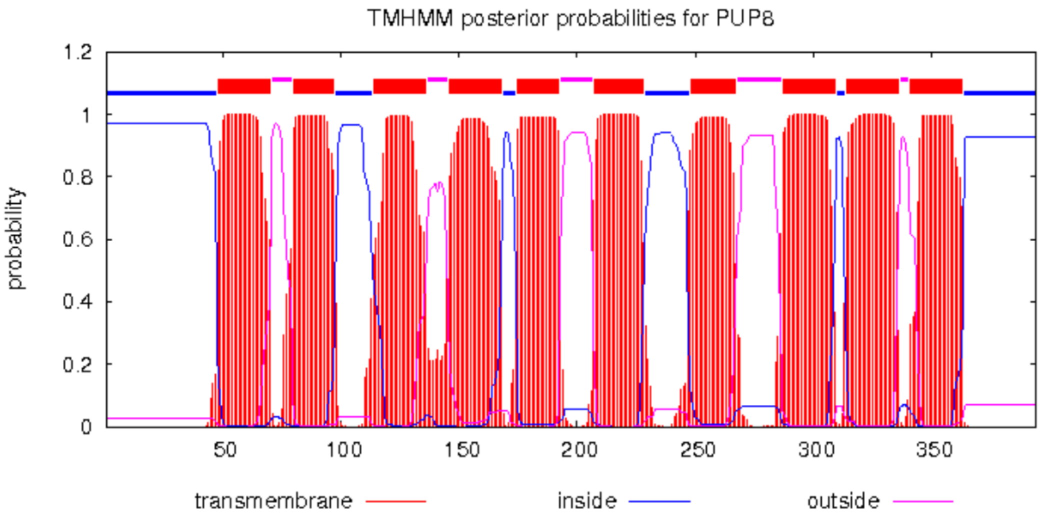


**Supplementary Figure S1.** Predicted transmembrane domains of PUP8.

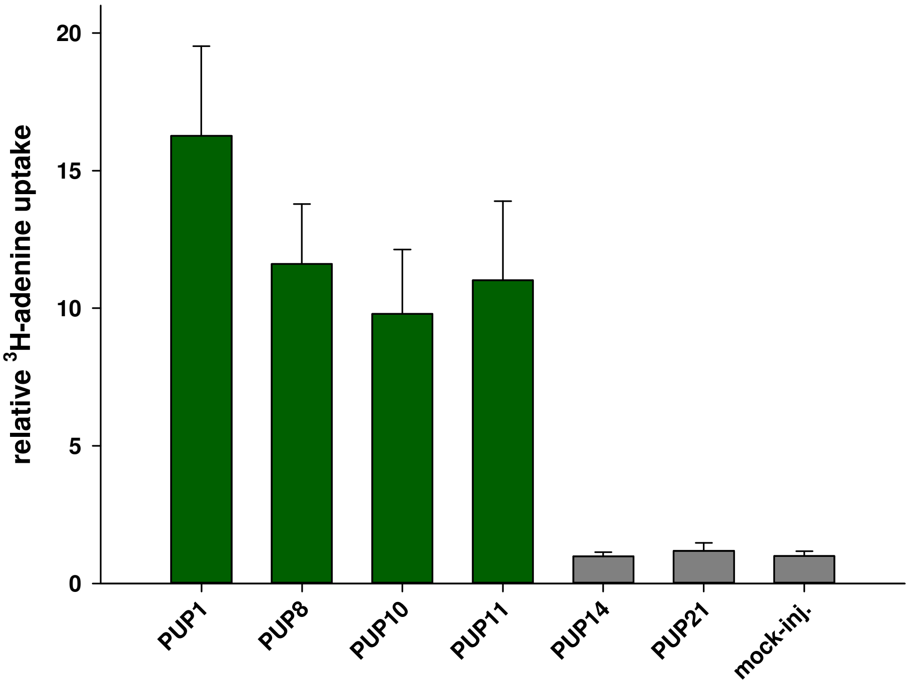


**Supplementary Figure S2. Adenine ($^3$H[adenine]) uptake assay in oocytes.** Oocytes expressing PUP genes were incubated in 1 µCi/ml $^3$H[adenine] (37.6 Ci/mmol) containing kulori buffer (pH 5.0) for 10 min. Radioactivity from single oocyte was quantified using liquid scintillation counting. Error bars represent ± s.d. of mean, n = 5-7 oocytes.

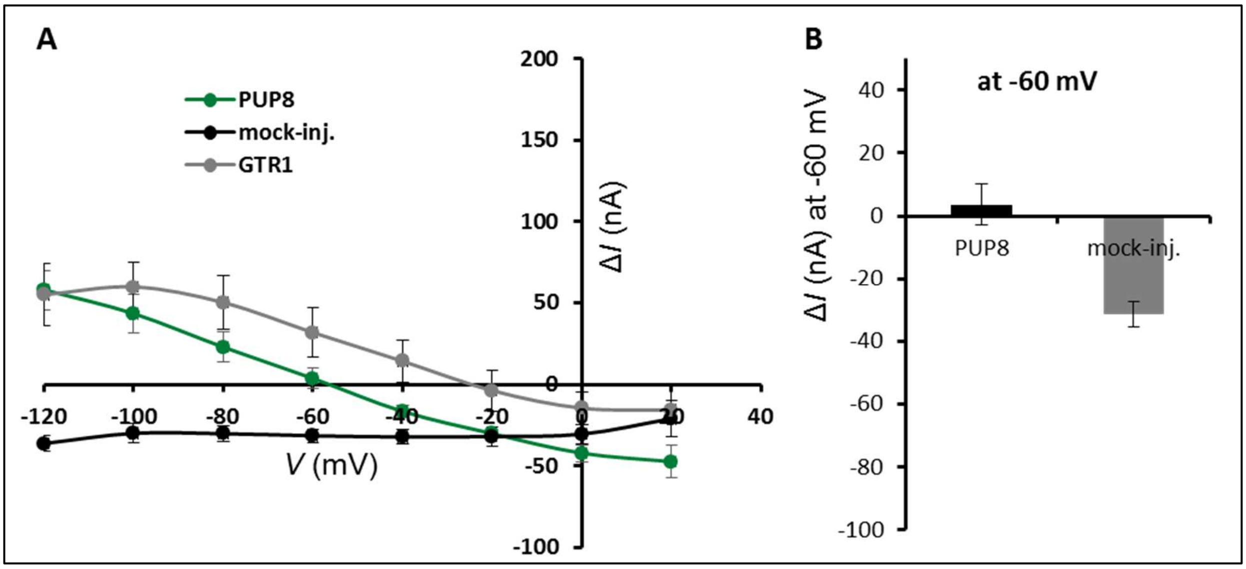


**Supplementary Figure S3. Electrophysiological characterization of PUP8-mediated phlorizin uptake.** PUP8-mediated phlorizin-induced currents were measured using the two-electrode voltage-clamp technique in a Robocyte. PUP8 or GTR1-expressing or mock-injected control oocytes were clamped to 8 different membrane potentials stepwise from -120 mV to +20 mV in 20 mV increments before and after addition of phlorizin (2 mM) in kulori buffer, and oocyte currents were measured under continuous perfusion of MES-based kulori buffer (pH 5) containing no or 2 mM phlorizin. **(A)** *IV* curve in response to phlorizin in eKulori (pH 5). **(B)** Bar graph of the average current induced in the presence of phlorizin at -60 mV at pH 5. Error bars represent ± s.d. of mean, n = 4 - 5 oocytes.

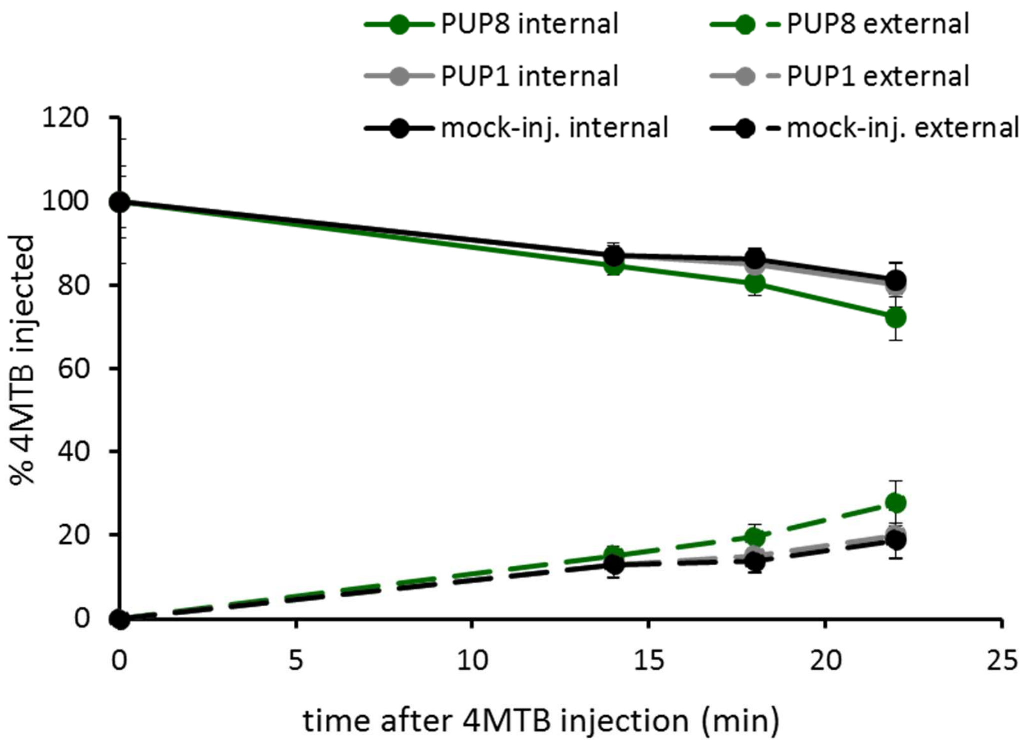


**Supplementary Figure S4. Injection-based 4MTB export assay in *Xenopus* oocytes.** Oocytes expressing PUP8, PUP1, or mock-injected oocytes were injected with 4MTB (to obtain initial internal concentration ~0.5 mM) and incubated in a kulori buffer (pH 7.4, without phlorizin). Export transport activity was measured by quantifying intracellular and extracellular phlorizin content, after 14, 18 and 22 h of incubation, using LC-MS/MS analysis. Error bars represent ± s.d. of mean of % 4MTB injected, n = 5 (5 x 3 oocytes).

**Supplementary Table S1.** List of primers used for *in vitro* transcription.

| **Primer name** | **Sequence (5' to 3' direction)** | **Reference** |
|---|---|---|
| HHN49-fw | AATTAACCCTCACTAAAGGGTTGTAATACGACTCACTATAGGG | (Wulff et al., 2019) |
| HHN50-rev | TTTTTTTTTTTTTTTTTTTTTTTTTTTTTTTATACTCAAGCTAGCCTCGAG | (Wulff et al., 2019) |
| Bolar051 FW1 | TGTGCTGAATTGTAATACGACTCACTATAGGGAGCTTGCTTGTTCTTTTTG C | (Larsen et al., 2017a) |
| Bolar051 RV2 | CCATTCGCCATTCAGGCT | (Larsen et al., 2017a) |

**Supplementary Table S2. List of strains used in this study**.

| Strain name | Genotype | Reference |
|---|---|---|
| BG | NCYC 3608 (*MATα hoΔ0 his3Δ0 leu2Δ0 ura3Δ0 cat5Δ0::CAT5(I91M) mip1Δ0::MIP1(A661T) gal2Δ0::GAL2 sal1Δ0::SAL1*) | (Eichenberger et al., 2017) |
| DBR2 | BG *[ARS/CEN/URA3/$P_{GPD1}$-HaCHS-$T_{CYC1}$/$P_{PGK1}$-ScTSC13-$T_{ADH2}$/$P_{TEF1}$-At4CL2-$T_{ENO2}$/$P_{PDC1}$-AtPAL2-$T_{FBA1}$/$P_{TEF2}$-AmC4H-$T_{PGI1}$/$P_{PYK1}$-ScCPR1-$T_{ADH1}$]* | (Eichenberger et al., 2017) |
| PHZ_PUP8 | DBR2 *[ARS/CEN/HIS3/$P_{GPD1}$- PcUGT88F2-$T_{CYC1}$/$P_{PGK1}$-PUP8-$T_{ADH2}$]* | this study |
| PHZ_PUP1 | DBR2 *[ARS/CEN/HIS3/$P_{GPD1}$- PcUGT88F2-$T_{CYC1}$/$P_{PGK1}$-PUP1-$T_{ADH2}$]* | this study |
| PHZ_control | DBR2 *[ARS/CEN/HIS3/$P_{GPD1}$- PcUGT88F2-$T_{CYC1}$/$P_{PGK1}$- stuffer -$T_{ADH2}$]* | this study |
| PUP8V | BG *[ARS/CEN/HIS3/$P_{GPD1}$- PcUGT88F2-$T_{CYC1}$/$P_{PGK1}$-PUP8-Venus-$T_{ADH2}$]* | this study |

**Supplementary Table S3.** List of primers used for cloning in this study.

| Primer name | Sequence (5' to 3' direction) | PCR products |
|---|---|---|
| **To construct pNB1u-PUP8** | | |
| ZM072_fw | GGCTTAAUGGCGCGCCATGCGCGGCTATG | D-tag-PGK1 |
| ZM0111_Rv | ATTTTAAGCUTTGTTTTATATTTGTTG | D-tag-PGK1 |
| ZM0112_fw | AGCTTAAAAUGGAAATAACTCAAGTAATC | PUP8 |
| ZM0113_rv | ATCCGCGGUCATACACTATGTATGTTTGTG | PUP8 |
| ZM0114_fw | ACCGCGGAUCTCTTATGTCTTTACGATTTATAG | ADH2-C-tag |
| ZM077_Rv | GGTTTAAUGGCGCGCCCGTGCCGTCGTTG | ADH2-C-tag |
| **To construct pNB1u-PUP1** | | |
| ZM072_fw | GGCTTAAUGGCGCGCCATGCGCGGCTATG | D-tag-PGK1 |
| ZM089_Rv | ATTTTAAGCUTTGTTTTATATTTGTTGTAAAAAG | D-tag-PGK1 |
| ZM086_Fw | AGCTTAAAAUGAAGAATGGTTTGATAATC | PUP1 |
| ZM087_Rv | AGATCCGCGGTUAAGCAACATAATCACTAAC | PUP1 |

| | | |
|---|---|---|
| ZM088_Fw | AACCGCGGATCUCTTATGTCTTTACGATTTATAG | ADH2-C-tag |
| ZM077_Rv | GGTTTAAUGGCGCGCCCGTGCCGTCGTTG | ADH2-C-tag |
| | **To construct pNB1u-PUP8V** | |
| ZM072_fw | GGCTTAAUGGCGCGCCATGCGCGGCTATG | D-tag-PGK1 |
| ZM0111_Rv | ATTTTAAGCUTTGTTTTATATTTGTTG | D-tag-PGK1 |
| ZM0112_fw | AGCTTAAAAUGGAAATAACTCAAGTAATC | PUP8 |
| ZM091_Rv | AGGTTTAAUCCTACACTATGTATGTTTGTG | PUP8 |
| ZM082_fw | ATTAAACCUCAGCCTGGGTAGCGGTGGAATG | Venus |
| ZM098_Rv | AGATCCGCGGUTACTTGTACAGCTCGTCCATGC | Venus |
| ZM084_fw | ACCGCGGATCUCTTATGTCTTTACGATTTATAG | ADH2-C-tag |
| ZM077_Rv | GGTTTAAUGGCGCGCCCGTGCCGTCGTTG | ADH2-C-tag |

**Supplementary Table S4. List of plasmids carrying homologous recombination tags used in this study.**

| Plasmid name | Content | **Reference** |
|---|---|---|
| pEVE2176 | *B-tag* $P_{GPD1}$- *PcUGT88F2*-$T_{CYC1}$ *C-tag* | (Eichenberger et al., 2017) |
| pEVE2177 | *C-tag* $P_{PGK1}$-*stuffer2*-$T_{ADH2}$ *D-tag* | (Eichenberger et al., 2017) |
| pNB1u-PUP8 | *C-tag* $P_{PGK1}$- *PUP8*-$T_{ADH2}$ *D-tag* | this study |
| pNB1u-PUP1 | *C-tag* $P_{PGK1}$- *PUP1*-$T_{ADH2}$ *D-tag* | this study |
| pNB1u-PUP8V | *C-tag* $P_{PGK1}$- *PUP8-Venus*-$T_{ADH2}$ *D-tag* | this study |
| pEVE1968 | *A-tag ARS/CEN CmR B-tag* | (Eichenberger et al., 2017) |
| pEVEDZ | *D-tag closing linker Z-tag* | (Eichenberger et al., 2017) |
| pEVEZA | *Z-tag pSC101 HIS3 A-tag* | (Eichenberger et al., 2017) |

| | | |
|---|---|---|
| pPHZPUP8 | *ARS/CEN/HIS3/$P_{GPD1}$- PcUGT88F2-$T_{CYC1}$/$P_{PGK1}$-PUP8-$T_{ADH2}$* | this study |
| pHZPUP1 | *ARS/CEN/HIS3/$P_{GPD1}$- PcUGT88F2-$T_{CYC1}$/$P_{PGK1}$-PUP1-$T_{ADH2}$* | this study |
| pPHZcontrol | *ARS/CEN/HIS3/$P_{GPD1}$- PcUGT88F2-$T_{CYC1}$/$P_{PGK1}$- stuffer-$T_{ADH2}$* | this study |
| pPUP8V | *ARS/CEN/HIS3/$P_{GPD1}$- PcUGT88F2-$T_{CYC1}$/$P_{PGK1}$-PUP8-Venus-$T_{ADH2}$* | this study |

**Supplementary Table S5.** MRM transitions for LC-MS/MS analysis.

| **Analyte** | **Retention Time [min]** | **Q1 [*m/z*]** | **Q3 [*m/z*]** | **CE [eV]** |
|---|---|---|---|---|
| Phlorizin | 1.79 | 435.1 | 273.0[Qt] | 13 |
| $[M-H]^-$ | | 435.1 | 167.0 | 27 |
| | | 435.1 | 123.1 | 39 |
| Sinigrin (IS) | 1.31 | 358.0 | 97.0[Qt] | 22 |
| $[M-H]^-$ | | 358.0 | 75.0 | 30 |
| | | 358.0 | 259.0 | 20 |
| 4MTB GLS | 1.74 | 420.0 | 97.0[Qt] | 23 |
| $[M-H]^-$ | | 420.0 | 259.0 | 23 |
| | | 420.0 | 75.0 | 30 |

Qt = quantifier ion, additional transitions are used for identification only. IS=internal standard; CE=collision energy; Q=quadrupole.
